# Transatlantic spread of highly pathogenic avian influenza H5N1 by wild birds from Europe to North America in 2021

**DOI:** 10.1101/2022.01.13.476155

**Authors:** V. Caliendo, N. S. Lewis, A. Pohlmann, S.R. Baillie, A.C. Banyard, M. Beer, I.H. Brown, R.A.M. Fouchier, R.D.E. Hansen, T.K. Lameris, A.S. Lang, S. Laurendeau, O. Lung, G. Robertson, H. van der Jeugd, T.N. Alkie, K. Thorup, M.L. van Toor, J. Waldenström, C. Yason, T. Kuiken, Y. Berhane

**Affiliations:** Department of Viroscience, Erasmus University Medical Center; Rotterdam, the Netherlands; Department of Pathobiology and Population Sciences, Royal Veterinary College; Hatfield, United Kingdom; Institute of Diagnostic Virology, Friedrich-Loeffler-Institut; Greifswald - Insel Riems, Germany; Animal and Plant Health Agency; Addlestone, United Kingdom; Vogeltrekstation - Dutch Centre for Avian Migration and Demography NIOO-KNAW; Wageningen, The Netherlands; Department of Biology, Memorial University of Newfoundland; St. John’s, Canada; Canadian Food Inspection Agency; Winnipeg, Canada; Environment and Climate Change Canada; Mount Pearl, Canada; Linnaeus University; Kalmar, Sweden; Atlantic Veterinary College, University of Prince Edward Island; Charlottetown, Canada; British Trust for Ornithology; Norfolk, United Kingdom; Natural History Museum of Denmark, University of Copenhagen; Copenhagen, Denmark; Center for Macroecology, Evolution and Climate, Globe Institute, University of Copenhagen; Copenhagen, Denmark; Department of Coastal Systems, Royal Netherlands Institute for Sea Research; Den Burg, the Netherlands; European Union for Bird Ringing c/o British Trust for Ornithology; Norfolk, United Kingdom

## Abstract

Highly pathogenic avian influenza (HPAI) viruses of the A/Goose/Guangdong/1/1996 lineage (GsGd), which threaten the health of poultry, wildlife and humans, are spreading across Asia, Europe and Africa, but are currently absent from Oceania and the Americas. In December 2021, H5N1 HPAI viruses were detected in poultry and a free-living gull in St. John’s, Newfoundland and Labrador, Canada. Phylogenetic analysis showed that these viruses were most closely related to HPAI GsGd viruses circulating in northwestern Europe in spring 2021. Analysis of wild bird migration suggested that these viruses may have been carried across the Atlantic via Iceland, Greenland/Arctic or pelagic routes. The here documented incursion of HPAI GsGd viruses into North America raises concern for further virus spread across the Americas by wild bird migration.

**One-Sentence Summary:** Detection of H5N1 highly pathogenic avian influenza in Canada raises concern for spread in the Americas by migratory birds.

## Main Text

The A/Goose/Guangdong/1/96 (GsGd) lineage of highly pathogenic avian influenza (HPAI) H5 virus first emerged in poultry in Southeast Asia more than 25 years ago. During the first decade of circulation of this lineage, the hemagglutinin (H) genes diversified into multiple genetic clades. GsGd viruses of clade 2.3.4.4 started to dominate outbreaks globally from 2014 onwards with clade 2.3.4.4b currently emerging as a particularly fit virus. This lineage, and particularly clade 2.3.4.4b, is expanding both its geographical spread and its host range (*1–3*). Therefore, this lineage of HPAI H5 virus is an increasing threat to the health of poultry, wildlife, and humans worldwide, as well as a growing economic problem for the global poultry sector.

In recent years, HPAI GsGd H5 outbreaks have frequently occurred in Europe (*3,4*). For the first time in 2005, the virus spread from Asia to Russia, western Europe, Africa and the Middle East, causing high mortality in wild birds and poultry (*5*). This spread was a result of unprecedented long-distance transport of HPAI virus, in which wild migratory ducks, geese and swans were implicated. Nine years later, in 2014, this happened again, and since the 2014/2015 outbreak, HPAI H5 viruses have caused ever larger outbreaks in wild birds and poultry in Europe nearly every year (*4*). In addition, there are also growing concerns about the zoonotic risks, and in December 2021, the European Centre for Disease Prevention and Control raised the risk level for virus transmission to occupationally exposed people from ‘low’ to ‘low/moderate’ (*4,6*).

In December 2021, there was a die-off of domestic birds on an exhibition farm in St. John’s, a city on the Avalon Peninsula of the island of Newfoundland, on the Atlantic coast of Canada (Supplementary Material). The cause was diagnosed as H5N1 HPAI. This was the first report of HPAI H5 in the Americas since June 2015, when the virus spread with wild birds across the Bering Strait to the Pacific coasts of Canada and the USA via the Pacific Flyway, one of the main avian migration routes (*7*). Genetic analysis showed the H gene corresponded to Eurasian HPAI viruses circulating in 2021 (*8*). This implied that the virus had been carried across the Atlantic, a route that had not been recorded before for any HPAI virus. Therefore, the goal of this study was 1) to investigate in detail whether the HPAI cases in Newfoundland were linked to the recent (2020/2021) or currently ongoing (2021/2022) HPAI outbreaks in Europe, and 2) to indicate the most likely scenario by which the virus crossed the Atlantic with migratory birds.

Phylogenetic analyses were performed to compare the genome sequences of the Newfoundland viruses from the exhibition farm birds and a great black-backed gull (Supplementary Table 1) found nearby to other influenza viruses in the database. Based on maximum likelihood and time-resolved trees inferred by use of whole genome sequences, the Newfoundland viruses had a shared common ancestor with European viruses circulating in early 2021 (Figure 1; Supplementary Figure 1). The dates for the most recent common ancestor (MRCA) of all gene segments ranged from December 2019 to April 2021 (Table 1). There was no evidence that the Newfoundland viruses were genetically closely related to other current or recent viruses circulating in Europe. In contrast to currently circulating European viruses, the sequences of the Newfoundland viruses had no evidence of reassortment with other avian influenza viruses after ancestral emergence (Supplementary Figure 2). The virus from the great black-backed gull was highly similar to those from the exhibition farm, except for a small number of nucleotide differences in the neuraminidase (N) gene segment.

**Fig. 1.**
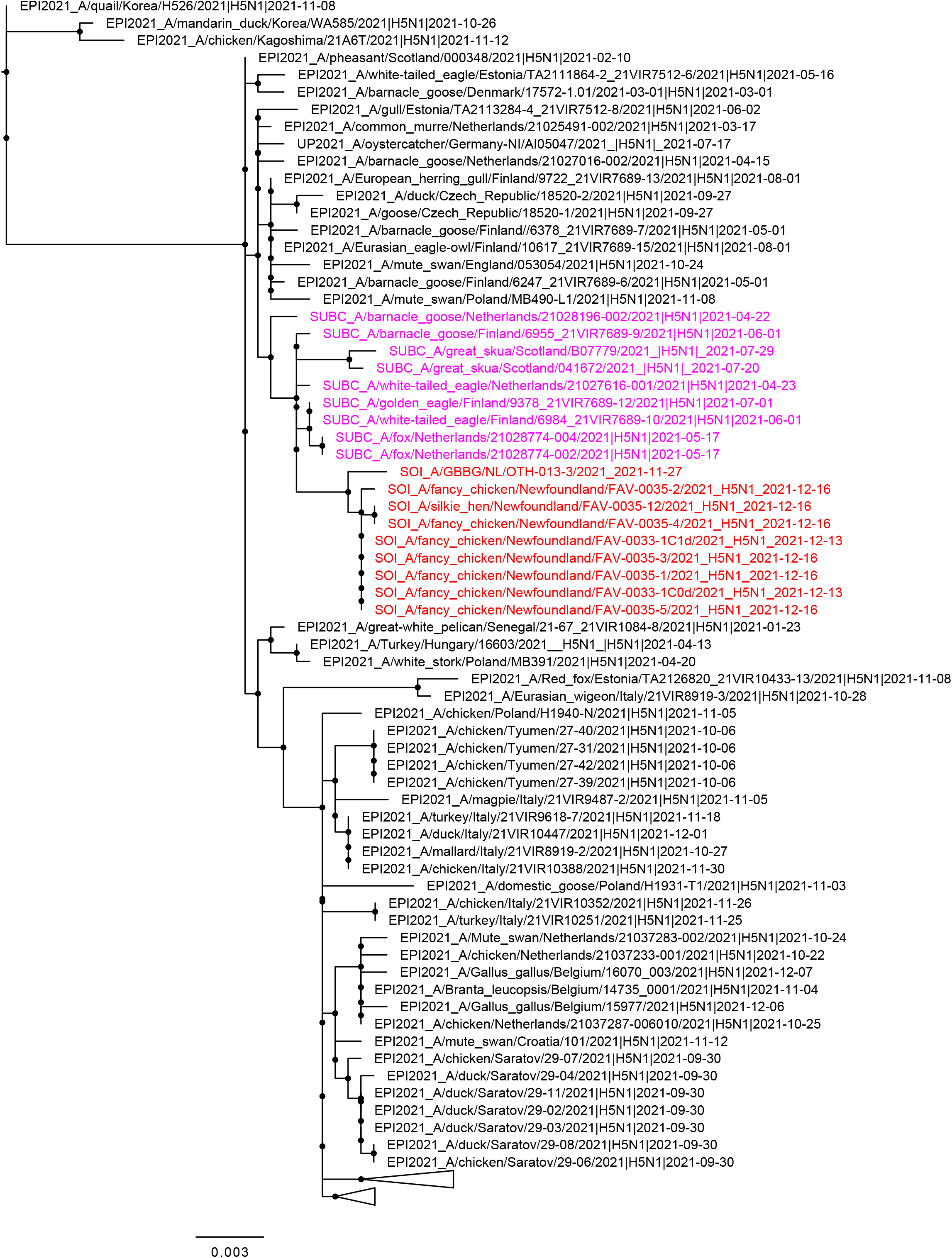
(because of its large size figure 1 is submitted as related document) Maximum likelihood phylogenetic tree of the H5 HA gene. Relationships among the European 2021 H5 2.3.4.4b HPAI strains (magenta) and the Newfoundland wild bird and outbreak strains (red) are shown. The tree is rooted by the outgroup and nodes placed in descending order. Clades are collapsed for clarity.

**Table 1.**
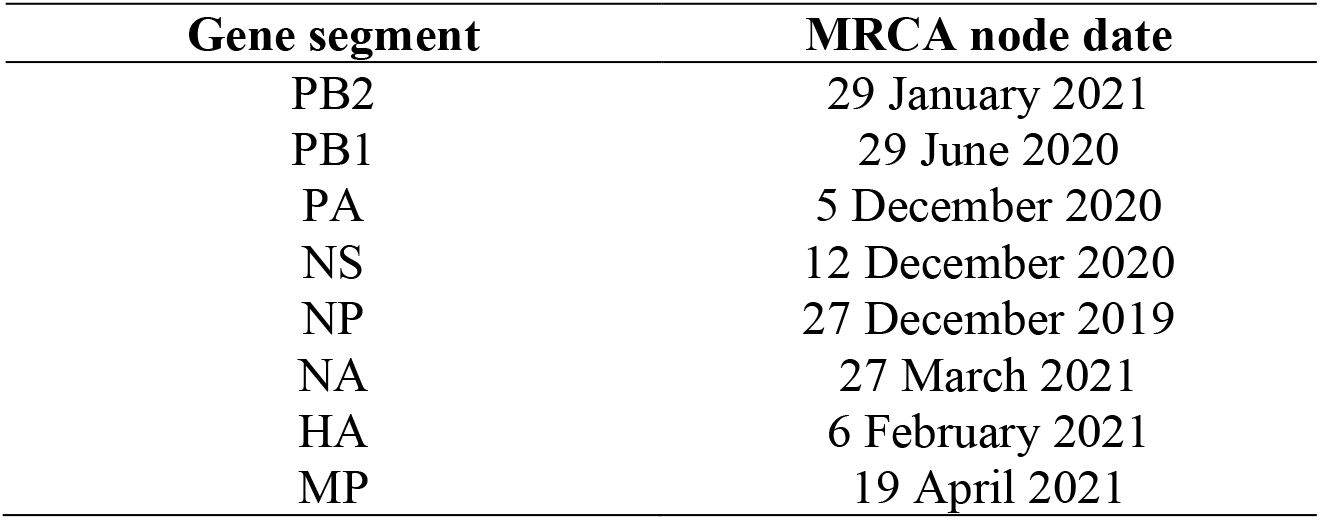
Dates for the most recent common ancestor (MRCA) of all gene segments.

There are several pathways for HPAI H5N1 virus to be carried across the Atlantic with migrating birds, based on the multitude of migration routes of wild birds and their overlapping ranges at breeding, stop-over, and wintering sites. Ring-recovery data confirm the regular movements of wild birds from Europe to Iceland and other North Atlantic islands, and from there to North America, and also provide evidence for direct movements of for example seabirds and gulls (Supplementary Table 2). Ringed individuals with a European origin have been found on Newfoundland for 8 of the 24 species in Supplementary Table 2. Given that the most likely ancestor of the virus detected in Newfoundland was circulating in Northwest Europe between October 2020 to April 2021 (see above), likely routes include spring migration of bird species moving to Icelandic, Greenlandic or Canadian High Arctic breeding grounds, or migration directly across the Atlantic Ocean (Figure 2; Supplementary Material).

**Fig. 2.**
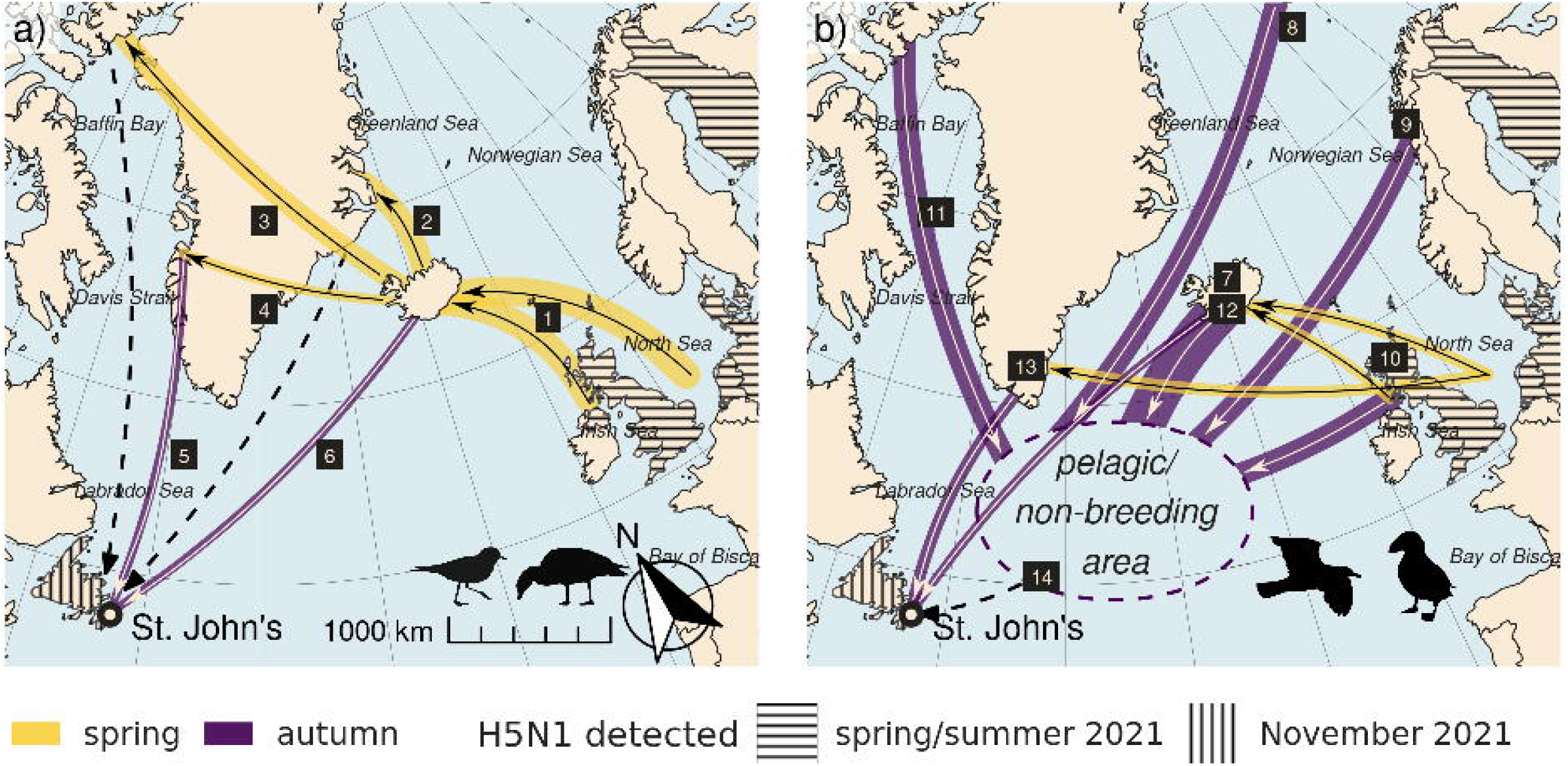
(because of its large size figure 2 is submitted as related document) Maps of transatlantic migration. Putative virus transmission pathways between Europe and Newfoundland via migratory waterfowl/shorebirds (a) and pelagic seabirds (b). Many Icelandic waterfowl and shorebirds (panel a) winter in Northwest Europe and return to Iceland to breed in spring (1), including whooper swans, greylag geese, pink-footed geese, Eurasian wigeons, Eurasian teals, northern pintails, common ringed plovers and purple sandpipers. Some bird populations use Iceland as a stopover site, and continue to breeding grounds in East Greenland (2; barnacle geese and pink-footed geese), the East Canadian Arctic (3; light-bellied brent geese, red knots, ruddy turnstones) and West Greenland (4; greater white-fronted geese). Migratory birds from Europe share these breeding areas with species that winter in North America, including Canada geese and snow geese from East Greenland and the East Canadian Arctic (5), and some Iceland-breeding species of duck, including small numbers of Eurasian wigeons, Eurasian teals, and tufted ducks (6). Several seabird species (panel b), such as gulls, skuas, fulmars and auks, have large breeding ranges in the Arctic. After the breeding season many species become fully pelagic and can roam large parts of the northern Atlantic. The mid-Atlantic ridge outside Newfoundland is an important non-breeding area for seabirds, and is frequented by auks from Iceland (7), Svalbard (8) and Norway (9), including large numbers of Atlantic puffins and Brünnich guillemots, and by black-legged kittiwakes and northern fulmars originating from Iceland, Norway and the United Kingdom (7-8, 10). There these birds are joined by seabirds from Canadian and Greenlandic waters (11). Direct migratory links to Newfoundland occurs through greater and lesser-black backed gulls as well as black-headed gulls from Iceland and Greenland (12, 13), and gulls also link the pelagic and the coastal zone around Newfoundland (14). Thickness of the lines highlights the relative approximate population sizes. Dashed lines show where small numbers of individuals, or vagrants, provide a potential pathway. For more details on species and population numbers see Supplementary Table 2.

Having reached the Avalon Peninsula of Newfoundland via one of above routes, the virus may have spread further within the abundant local populations of ducks and gulls wintering in the city of St. John’s. As these species are peridomestic and especially the ducks are known to feed around poultry farms, they may be candidates for incursion of the virus into the farm in St John’s.

To further evaluate which wild bird species might have been involved in transatlantic transport of HPAI H5 virus, we compared above bird migration patterns with reports of HPAI-H5-positive wild birds in Europe (Supplementary Table 3). We limited our evaluation to the period of six months up to April 2021, the latest MRCA date of the Newfoundland virus gene segments (see above), and to the coastal countries in Northwest Europe, which act as the main wintering areas for wild birds that migrate across the Atlantic. In that period and region of Europe, HPAI-H5-positive species included 15 species of ducks, geese, swans, gulls, and shorebirds. These species might be considered to be possible carriers of HPAI H5 virus from Europe in late winter 2020/2021 or early spring 2021 partly or all the way to Newfoundland (Supplementary Table 3). However, given the incompleteness of sampling and the possibility of wild birds carrying HPAI virus subclinically, the involvement of other wild bird species in transatlantic virus transport cannot be ruled out.

In conclusion, the HPAI H5N1 viruses that were detected in Newfoundland in November and December 2021 originated from Northwest Europe and belong to HPAI clade 2.3.4.4b._Most likely, these viruses emerged in Northwest Europe in winter 2020/2021, dispersed from Europe in late winter or early spring 2021, and arrived in Newfoundland in autumn 2021. The viruses may have been carried across the Atlantic by migratory birds using different routes, including Icelandic, Greenland/Arctic, and pelagic routes._The unusually high presence of the viruses in European wild bird populations in late winter and spring 2021, as well as the greater involvement of barnacle and greylag geese in the epidemiology of HPAI in Europe since October 2020, may explain why spread to Newfoundland happened this winter (2021/2022), and not in the previous winters.

The incursion of these HPAI viruses, which appear to be well-adapted to wild birds, raises concern about the potential of HPAI virus to become established and spread in the Americas via wild birds. If these viruses become established in the Atlantic Flyway, they could rapidly spread west to Mississippi, Central and Pacific Flyways, as has been seen for West Nile virus following its emergence in New York in 1999 (*9*). The implication of this scenario would be high wild bird mortality, risk for incursion into poultry holdings and those of other captive birds, as well as zoonotic risk.

To prevent and mitigate the risk of viral spread, it will be vital to increase surveillance of wild birds in eastern North America, as well as at migration stop-over stations in Iceland and Greenland. This should include virus detection with whole genome sequencing to enable molecular epidemiology. Collecting wild bird mortality reports can give an idea of the impact of the outbreak on local wild bird populations, and active surveillance is critical to identify vector species. The overlap of migratory movements of wild waterbirds along the Atlantic coast of North America with densely populated poultry areas may increase the risk of viral incursion into poultry farms, emphasizing the need for appropriate biosecurity measures and spatial planning of the poultry sector. The spread of HPAI H5 viruses from Europe to Canada stresses the importance of close international cooperation and data exchange to better understand the global epidemiology of avian influenza, and is a call to re-assess the poultry sector in a way that embraces the One Health perspective: to sustainably balance and optimize the health of people, animals and ecosystems (https://www.who.int/groups/one-health-high-level-expert-panel).

## Acknowledgments

We thank A. Byrne, J. James, S. Reid, M.E.B. Jones, C. Pekarik, T. Hisanaga, W. Xu, J. Koziuk, M. Fisher, A. T. Mjøs,G. Gargallo, G.A. Gudmundsson, J.J. Madsen, D.Moss, S. Lopes for technical support, data generation, interpretation and discussions. We gratefully acknowledge the originating laboratories, where specimens were first obtained, and the submitting laboratories, where sequence data were generated and submitted to the EpiFlu Database of the Global Initiative on Sharing All Influenza Data (GISAID), on which part of this research is based. All contributors of data may be contacted directly via the GISAID website (http://platform.gisaid.org). European bird ringing schemes provided ring recovery data via the EURING databank..

## Author contributions

Conceptualization: RF, TK, YB, Methodology: NSL, AP, YB, Investigation: VC, NSL, AP, SRB, AB, RDEH, TKL, ASL, SL, OL, KT, GR, HJ, TNA, MLT, JW, CY, YB, Visualization: VC, NSL, AP, TKL, MLT, JW, Supervision: MB, IB, RF, GR, HJ, TK, YB, Writing - original draft: VC, NSL, AP, TKL, ASL, SL, GR, TK, YB, Writing - review & editing: VC, NSL, AP, SRB, AB, MB, IB, RF, RDEH, TKL, ASL, SL, OL, KT, GR, HJ, TNA, MLT, JW, CY, TK, YB.

## Data and materials availability

All data are available in the main text or the supplementary materials, and in the GISAID database (http://platform.gisaid.org).

## Supplementary Materials

Supplementary Text

Materials and Methods

Figs. S1 to S2

Tables S1 to S4

References (*10–47*)

## Supplementary Materials for

### Epidemiological description of outbreak on exhibition farm

The index farm where highly pathogenic avian influenza (HPAI) H5 virus in captive birds occurred was an exhibition farm in St. John’s, a city located on the Avalon Peninsula of the island of Newfoundland, Canada. The farm housed 409 birds of different species, namely chickens, guineafowl, peafowl, emus, domestic ducks, domestic geese, and domestic turkeys. On 9^th^ December, the farm owner first noticed mortality. On 13^th^ December, the farm owner reported the increased mortality to a local veterinarian. Autopsies on four chickens showed swollen heads and cutaneous haemorrhages. Oropharyngeal and cloacal swabs from these chickens tested positive for H5 avian influenza virus at the Atlantic Veterinary College, University of Prince Edward Island, and the Canadian Food Inspection Agency (CFIA) was notified. On 16^th^ December, by which time 306 birds (mostly chickens, turkeys and guineafowl) had died, staff of the CFIA collected tissue samples from dead chickens, as well as oropharyngeal and cloacal swabs from different species of captive birds still present (Supplementary Table 4), after which all remaining captive birds were culled. On 20^th^ December, the CFIA confirmed the diagnosis of HPAI of the subtype H5N1.

There was ample opportunity on the farm for mingling of captive and wild birds. Captive birds except emus were allowed to move freely in and out of the open pens in which they were housed. At an on-site pond, domestic ducks were reported to mingle with free-living mallards (Supplementary Table 1), and a snowy egret had been observed around 2^nd^ to 6^th^ December. Other wild birds reported on the farm were common starlings, feral pigeons, and unspecified gulls.

Retrospectively, HPAI H5N1 virus also was identified in tissues of a great black-backed gull found at a nearby pond in St. John’s. The gull had been found ill on 26^th^ November 2021 and taken to a local wildlife rehabilitation centre, where it died the following day.

### Potential routes for HPAI H5 virus to be carried across the Atlantic with migrating birds

The first possible route via Iceland seems to be the strongest link between Newfoundland and Europe (*10,11*), because it is a meeting point of breeding bird populations which winter in Europe as well as along the East coast of North America. Numerous species, totaling almost two million individual birds, migrate annually from northwestern Europe to breeding grounds in Iceland and beyond.,. Several populations breed on Iceland, including swans (whooper swan), geese (greylag goose, pink-footed goose), ducks (Eurasian wigeon, Eurasian teal, Northern pintail), gulls (great- and lesser black-backed gull, black-headed gull, black-legged kittiwake) and shorebirds (common ringed plover, purple sandpiper, Supplementary Table 2). In addition, several species (e.g. barnacle geese and pink-footed geese) migrating to breeding grounds further away (Greenland and/or Canadian High Arctic) make spring and autumn stopovers in Iceland, (*12,13*). This creates potential for the virus to have been spread northwards to Iceland (or further northward) in spring, where it could have circulated among breeding birds, or transmitted during autumn migration by species returning from the Arctic. Several Iceland-breeding species of ducks (Eurasian wigeon, Eurasian teal, tufted duck), gulls (lesser black-backed gull, blacklegged kittiwake, black-headed gull) and alcids (Brunnich’s guillemot, Atlantic puffin) winter along the Atlantic coast of North America in variable numbers (Supplementary Table 2). If the virus was transmitted to one of these populations during their stay on Iceland, it could have been spread to Newfoundland during the subsequent autumn migration. Importantly, Iceland-breeding Eurasian wigeons or Eurasian teals could be responsible for both the journey to Iceland from European wintering grounds, as well as the journey from Iceland to Newfoundland, where these species are frequently encountered as vagrants (Supplementary Table 2, *14, 15*).

The second possible route is via species that migrate from northwestern Europe to the Canadian High Arctic and/or Northwest Greenland. These include shorebirds (e.g. ruddy turnstone, red knot) and some geese (light-bellied brent goose and greater white-fronted goose). If the virus circulated in these breeding populations and then moved to other coastal marine bird populations bordering Baffin Bay, which include huge numbers of colonial seabirds and marine waterfowl (16, *17*), the virus could have followed a coastal or even pelagic route south with the large autumn migration of Arctic marine birds (various sea ducks, auks and larids, *18-20*) to emerge in Newfoundland. Alternatively, shorebirds and waterfowl may have played a role: several European-wintering populations have overlapping breeding grounds with populations wintering along the east coast of North America. Regarding geese, greater white-fronted geese share breeding grounds in western Greenland with Canada geese (*21*), which migrate south along the Canadian Atlantic coast. Also, brent geese have overlapping breeding grounds with snow geese (*12*). In addition, individual barnacle geese and pink-footed geese breeding in Greenland could also have travelled south to Newfoundland carrying the virus, as these birds are regular vagrants to the North American Atlantic coast (*22*). While geese occur only in small numbers on Newfoundland, two barnacle geese and four pink-footed geese, probably originating from Greenland breeding grounds, were observed in the autumn of 2021. St. John’s is the first major population center on a coastal route south from Baffin Bay/Davis Strait and along the Labrador Shelf, so emergence in eastern Newfoundland is consistent with this route.

Three wild bird species involved in the Iceland and/or Greenland/High Canadian Arctic routes deserve particular attention. Eurasian wigeon have been prominently involved in outbreaks in Eurasia, and are considered prime candidates for carrying HPAI virus over long distances (*23*). Also, during the first stages of an outbreak they are one of the first species to be detected HPAI virus positive, often without clinical signs. Barnacle geese and greylag geese, which congregate in Iceland, were in the top three most abundant species detected H5-positive in Europe in late winter and early spring 2021 (*4*). Given that both greylag and barnacle geese have populations breeding on Iceland/Greenland and wintering in Europe (particularly the UK), these two species are high on the list of probable vectors that transported the virus to Iceland/Greenland and finally to Newfoundland. The high involvement of infected geese in the HPAI dynamics, which was not seen before October 2020, together with the unusually high levels of HPAI H5 virus presence in wild birds in Northwest Europe in spring 2021, might also explain why HPAI H5 virus spread to Newfoundland this winter (2021/2022), and not in the previous winters (2020/2021, 2016/2017, 2014/2015, 2005/2006). It is, however, striking that no cases of HPAI H5 virus have been recorded on Iceland in 2021.

A third possible, pelagic, route is directly across the Atlantic Ocean. Such a route could have started with coastal and pelagic seabirds in Northwest Europe, where the virus may have remained undetected for much of the summer of 2021. A subsequent migration of seabirds to Newfoundland waters in the autumn of 2021 could have brought the virus to North America. The large populations of black-legged kittiwakes and northern fulmars that breed in the U.K. have long been known to frequent Newfoundland waters (*24*), and these movements have been corroborated by recent telemetry studies (*25*). Further, recent telemetry information reveals that millions of pelagic seabirds breeding all across the Atlantic congregate over the Mid-Atlantic Ridge in the central North Atlantic at all times of year (*26*), making a pelagic transmission route a possibility. From the pelagic wintering grounds off Newfoundland, a species that uses both pelagic and coastal habitats, possibly a gull, may have brought the virus to shore in St. John’s. Trans-Atlantic transmission via seabirds has been suggested for LPAI viruses, including detection of mosaic Eurasian-North American viruses in gulls and alcids (*27,28*).

### Materials and Methods

#### Phylogenetic analysis

Methods for phylogenetic analysis were the same as Sagulenko 2018 and Poen 2019 (*29, 30*). Full genome sequences were obtained from nine clinical or postmortem samples of captive birds at the exhibition farm, and from one postmortem sample of a great black-backed gull from a nearby city pond (Supplementary Table 1).

Sequences: We searched for H5NX whole genome sequences in GISAID from Europe, Asia and Africa where samples were collected from 01-01-2021 through 27-12-2021. To these existing data we added eight unpublished sequences from Newfoundland, and three additional unpublished sequences from European wild birds collected in the timeframe to the GISAID database.

We aligned the sequences using MAFFT v7.407 and trimmed to the starting ATG and ending STOP codon. Maximum-likelihood trees were inferred using IQ-TREE 2.1.3. and 1000 replicates for SH approximate likelihood ratio test. We used TreeTime, a Python-based framework for phylodynamic analysis using an approximate Maximum Likelihood approach to estimate ancestral states, and reroot trees to maximize temporal signals.

#### Analysis of avian migration

We evaluated the possible routes along which wild birds can migrate from Europe to North America, based on knowledge on existing migration routes as well as the retrieval of identification (bird) rings. We compared the information with the data of HPAI H5 virus-positive birds from Northwest European countries (i.e., UK, Ireland, Norway, Finland, Denmark, Germany, Netherlands, Belgium, France) that are the starting points, or are situated along these migratory routes. For the analysis, we prioritized the most abundant bird species, that also most frequently tested H5-positive during the 2020/2021 outbreak in Europe.

We focused on bird species susceptible to avian influenza (waterfowl, gulls, shorebirds and seabirds) which either bred or made a migratory stopover on Iceland, this being the most likely connection between Europe and Newfoundland. We identified wintering grounds, staging sites and breeding grounds based on literature, using mostly the database of Birds of the World (*31, 32*). We estimated the population sizes breeding in Iceland or passing through Iceland during migration using Fox & Leafloor (2018), Icelandic Institute of Natural History (2021) and van Roomen (2018) (*32–34*).

We calculated the number of individuals observed in Newfoundland from Ebird data (*13*). We extracted all observations from complete lists done between September – December, 2011 to 2021 on Newfoundland. For rare species (with less than 10 records annually) we also included sightings from incomplete lists. For every year and species, we calculated the maximum number of observed individuals per location, and added these to calculate the total number of individuals observed in Newfoundland for every year. We then calculated the average number of individuals observed annually between 2011 - 2021, and the number observed in 2021. We identified the most likely origin of birds encountered in Newfoundland using the database Birds of the World (*31*) as well as several other resources.

Ring-recovery data were obtained from the EURING Migration mapping Tool MMT, an online tool under development, that provides information on movements of ringed birds between pre-set areas within Europe and to other areas of the world, based on the EURING databank. These data were augmented with published (individuals recovered up to 2002, Lyngs 2003, *48-50*) and unpublished data (to 2011) of birds ringed in Greenland supplied by Copenhagen Bird Ringing Centre. All records of individual birds moving between Northwest Europe (Norway & Sweden, Germany & Denmark, Belgium & Netherlands, Great Britain & Ireland) *and* Iceland and Faroe islands, *or* Svalbard and other North Atlantic islands *or* Greenland, and individual birds moving between these areas *and* Canada *or* USA were selected. Prior to selection, unlikely records (finding date before ringing date, finding or ringing location not accurate etc.) were removed. For species not considered in the Migration Mapping Tool, only records of birds moving between Northwest Europe and Greenland, Canada or USA were available.

**Fig. S1.**
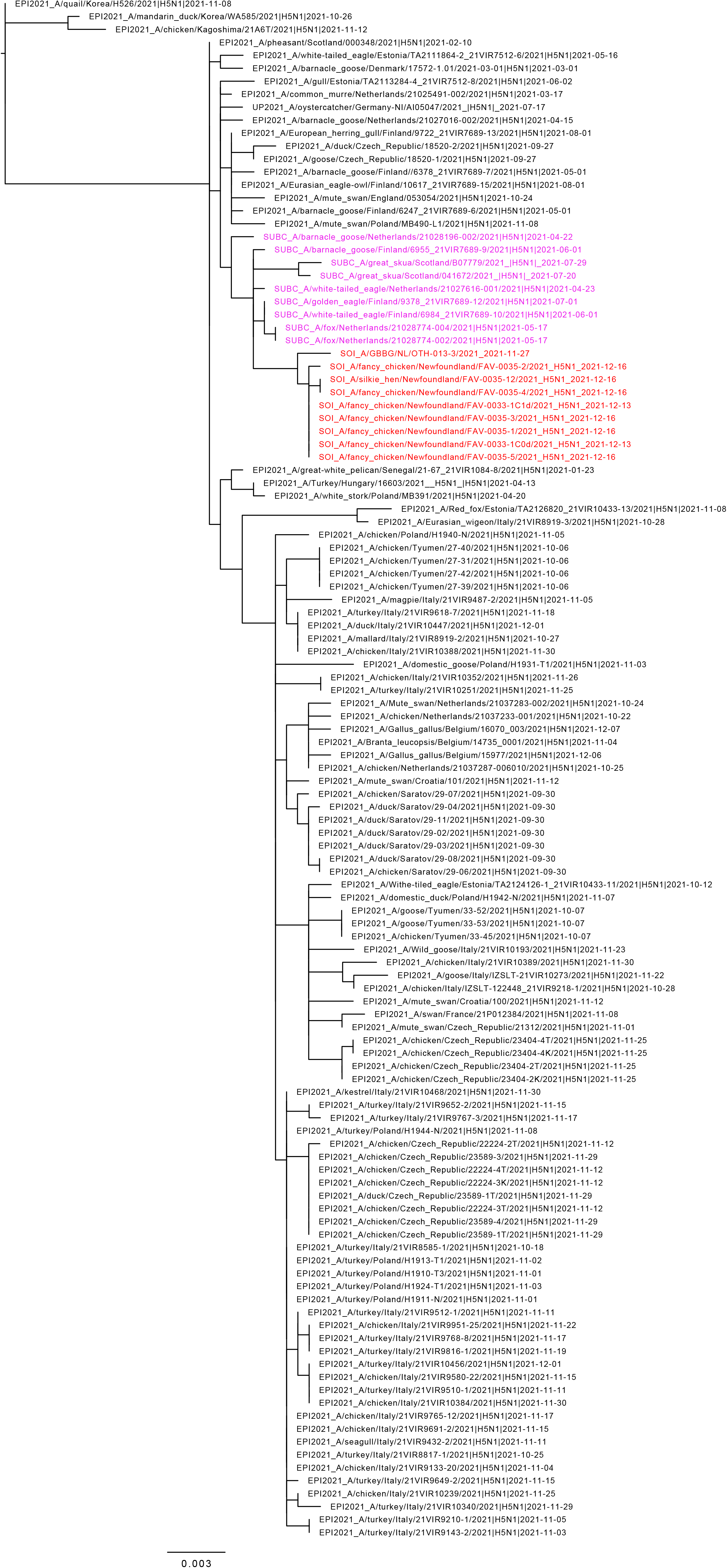
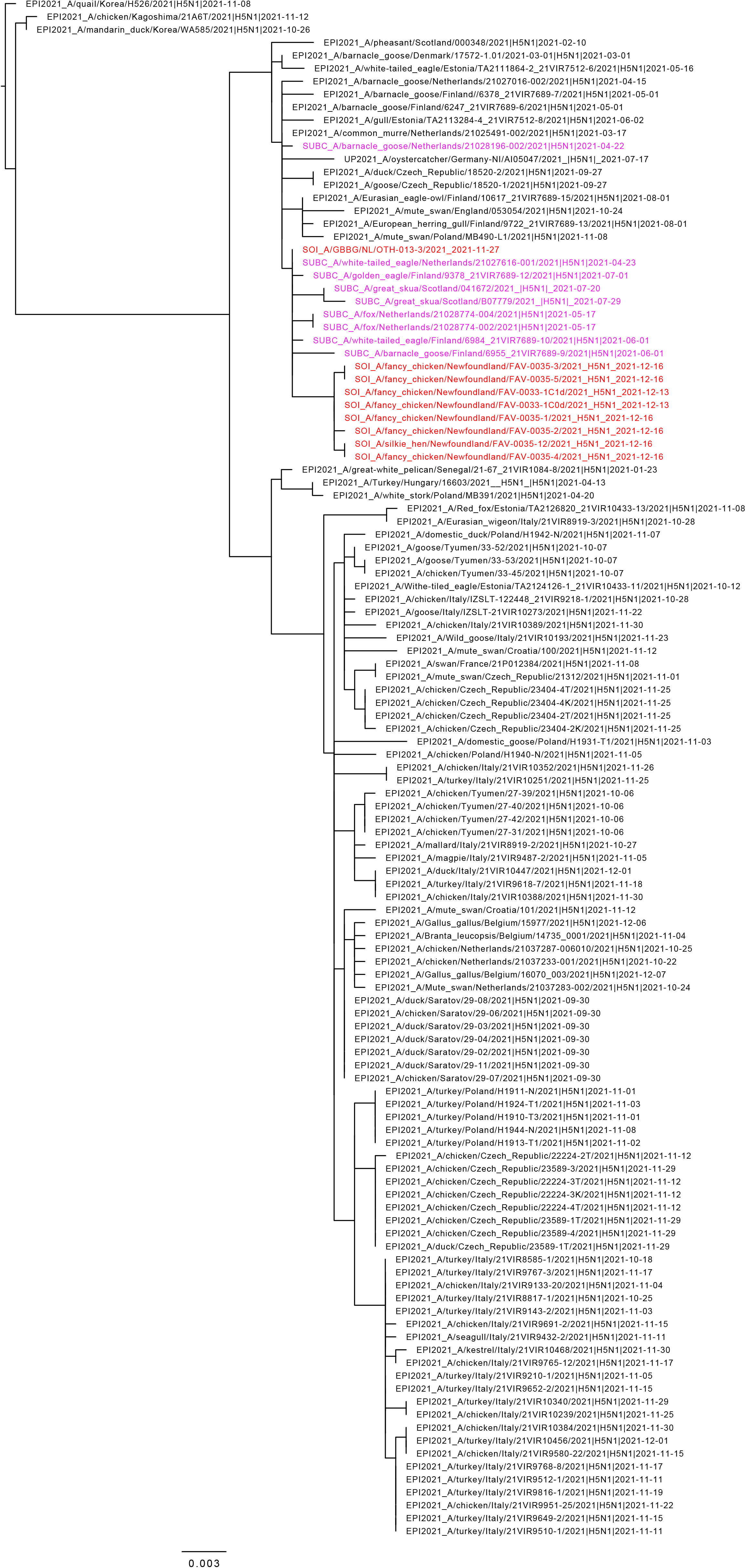
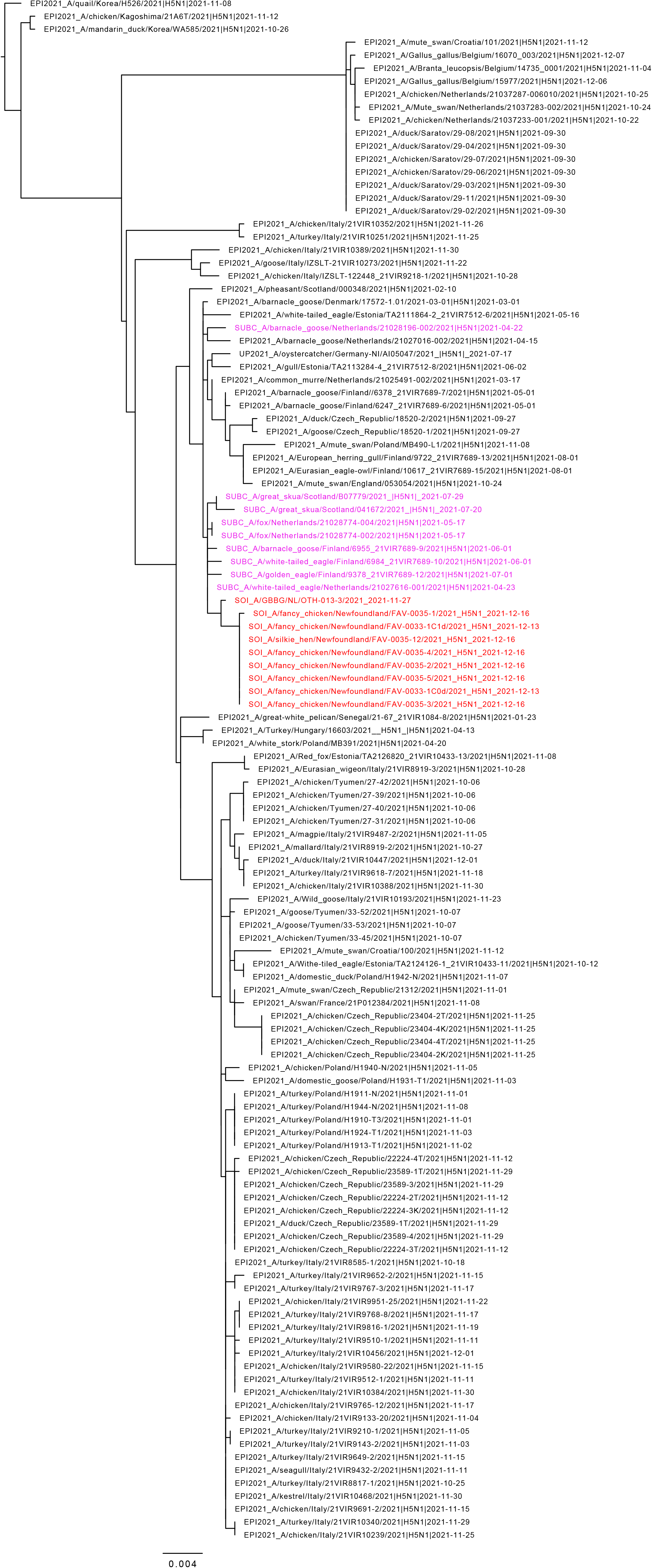
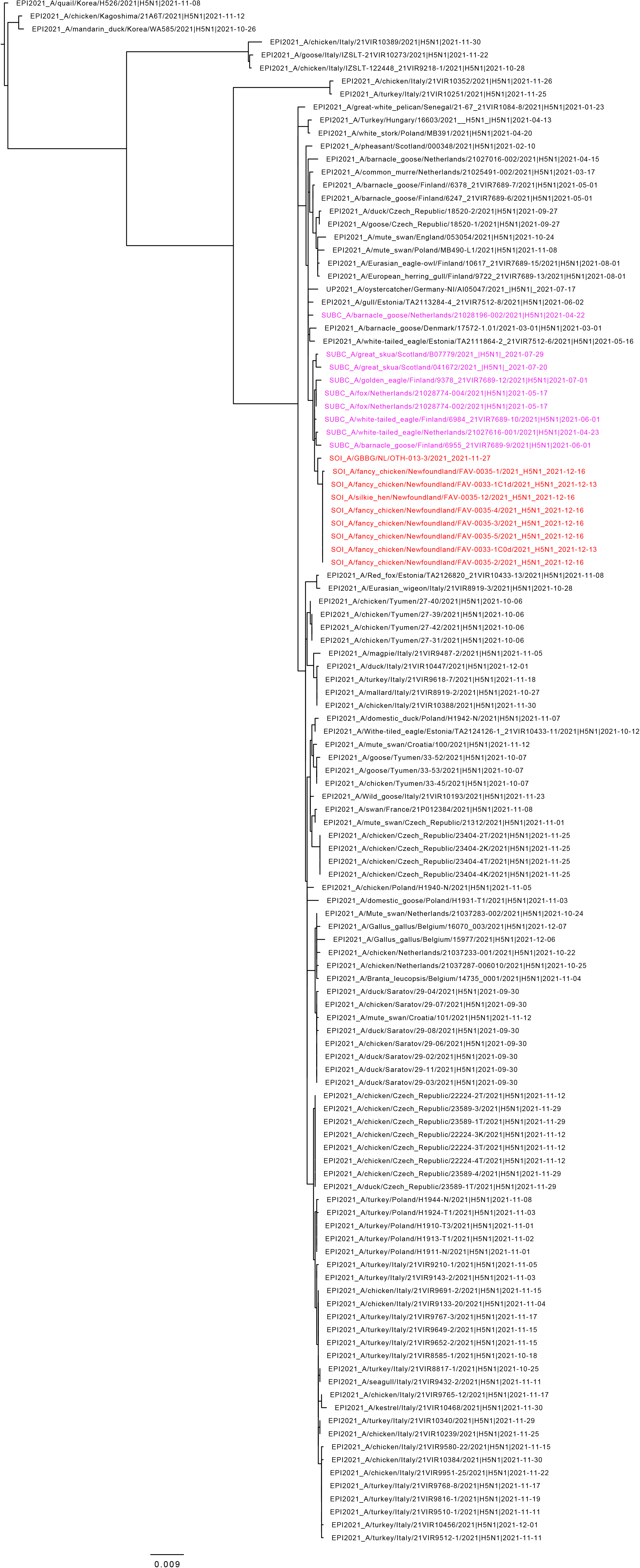
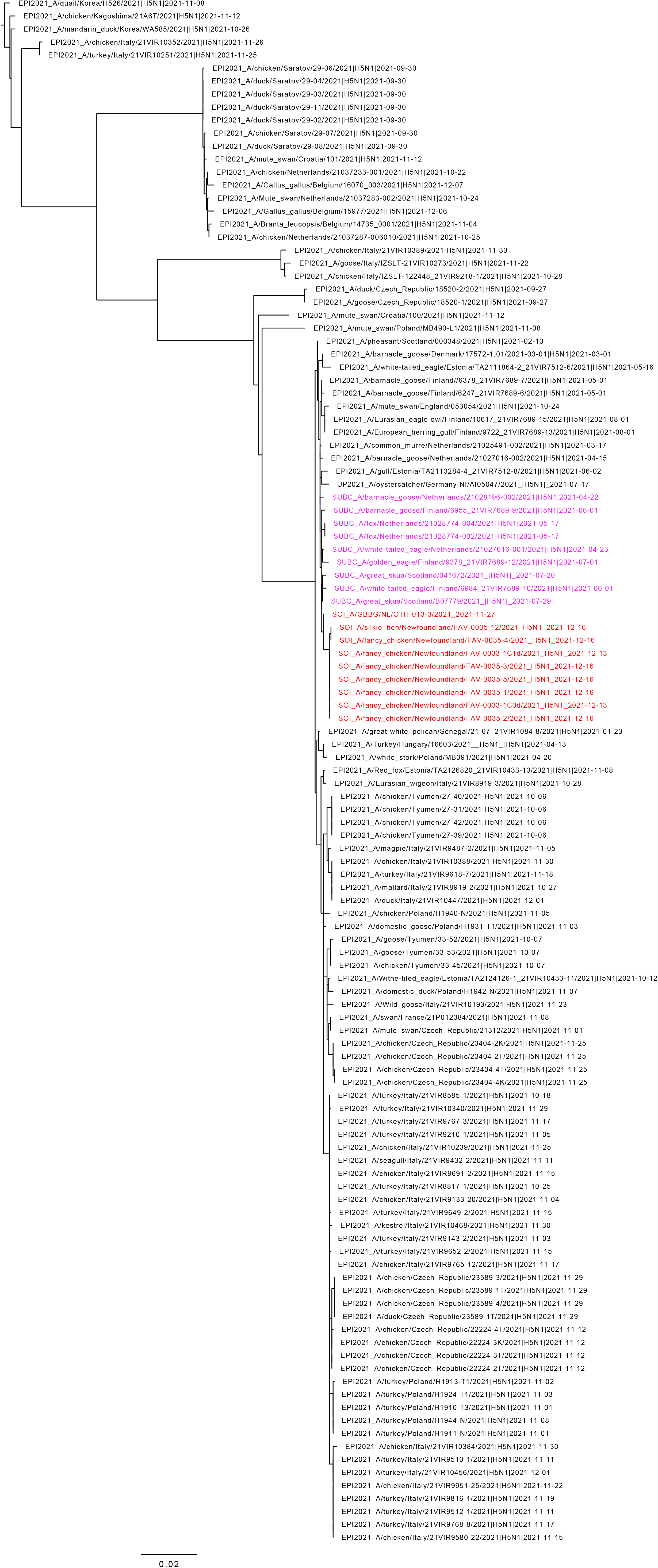
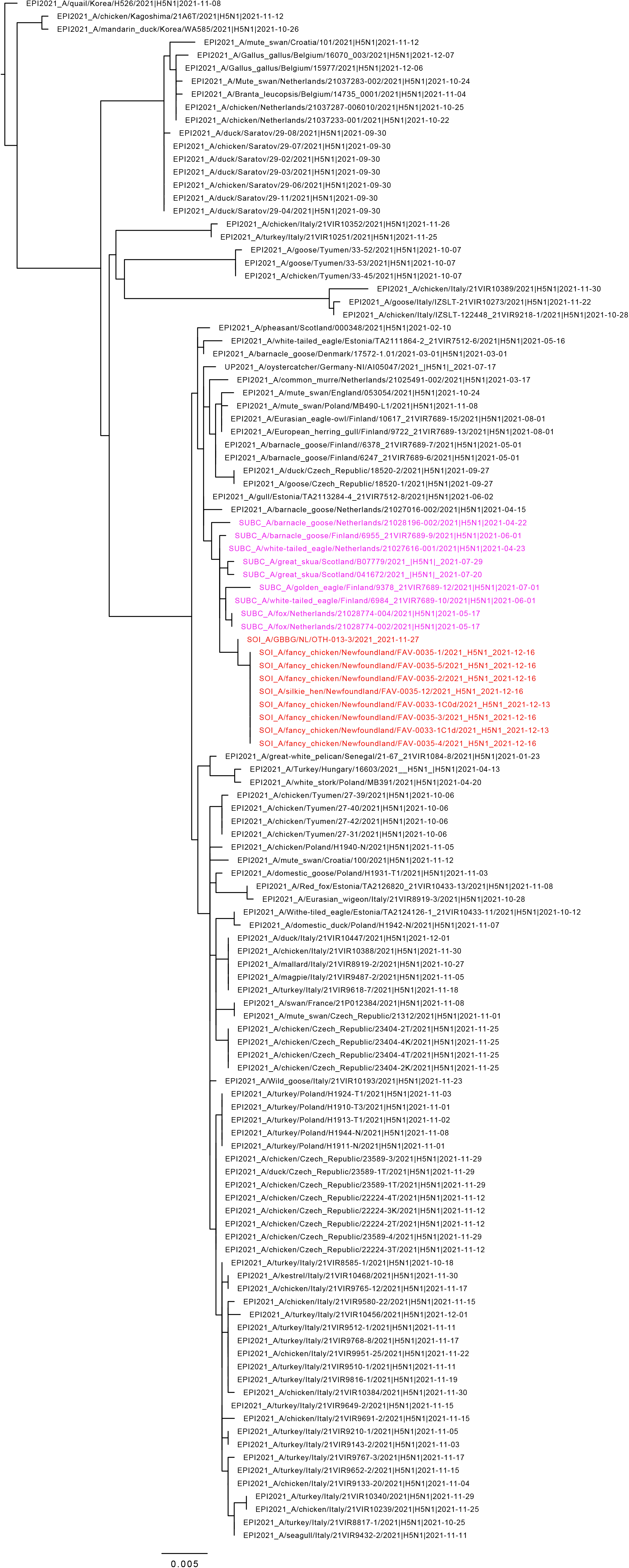
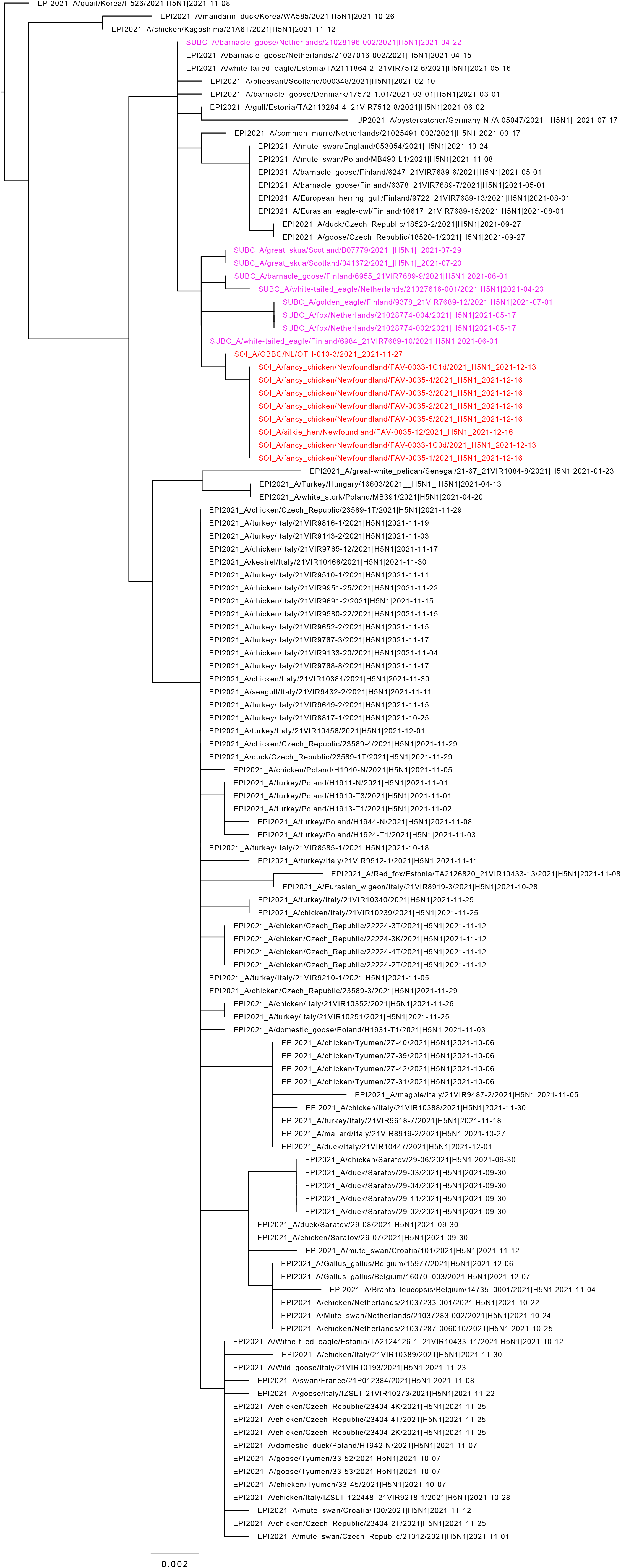
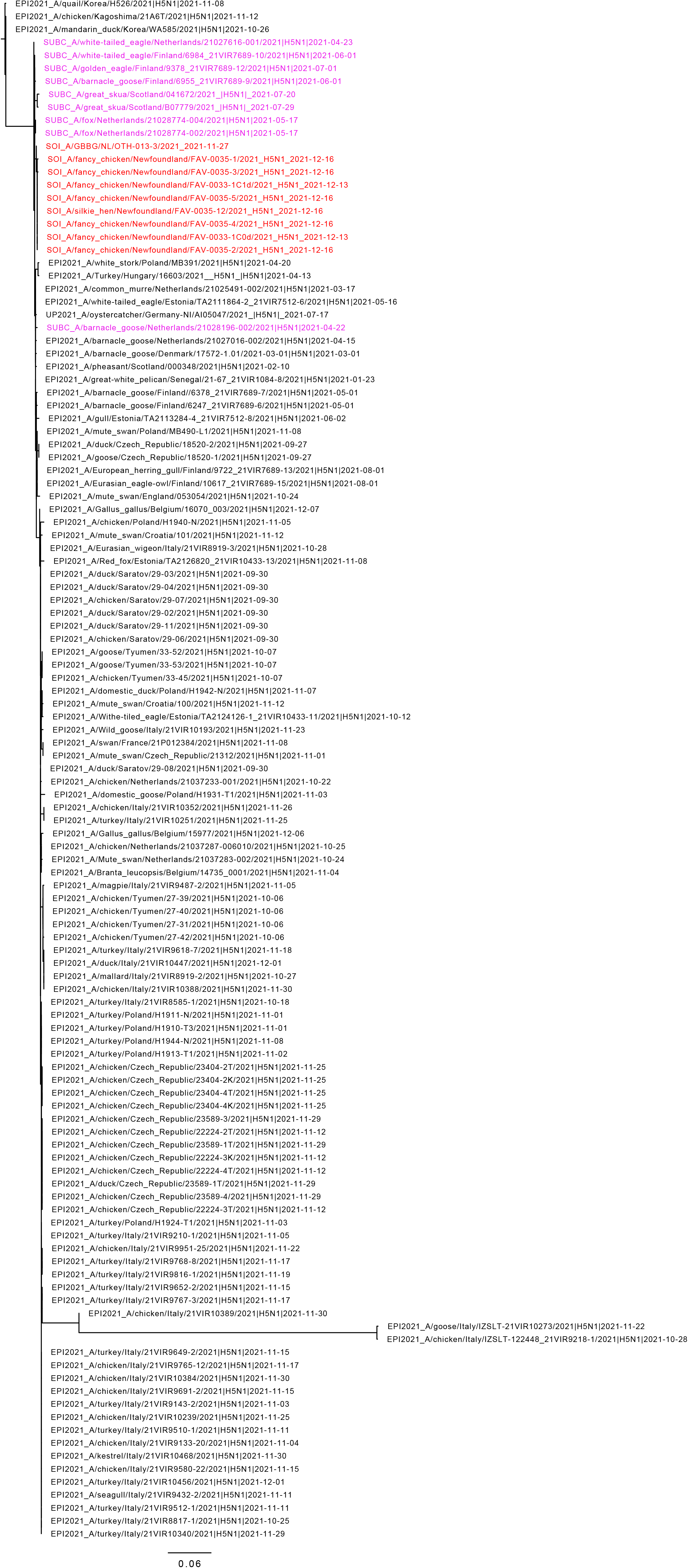
(because of its large size supplementary figure 1 is submitted as related document) Maximum likelihood phylogenetic tree of the H5 gene segments. Relationships among the European 2021 H5 2.3.4.4b HPAI strains (magenta) and the Newfoundland wild bird and outbreak strains (red) are shown. The tree is rooted by the outgroup and nodes placed in descending order; order: HA, NA, PA, PB1, PB2, NP, MP, NS.

**Fig. S2.**
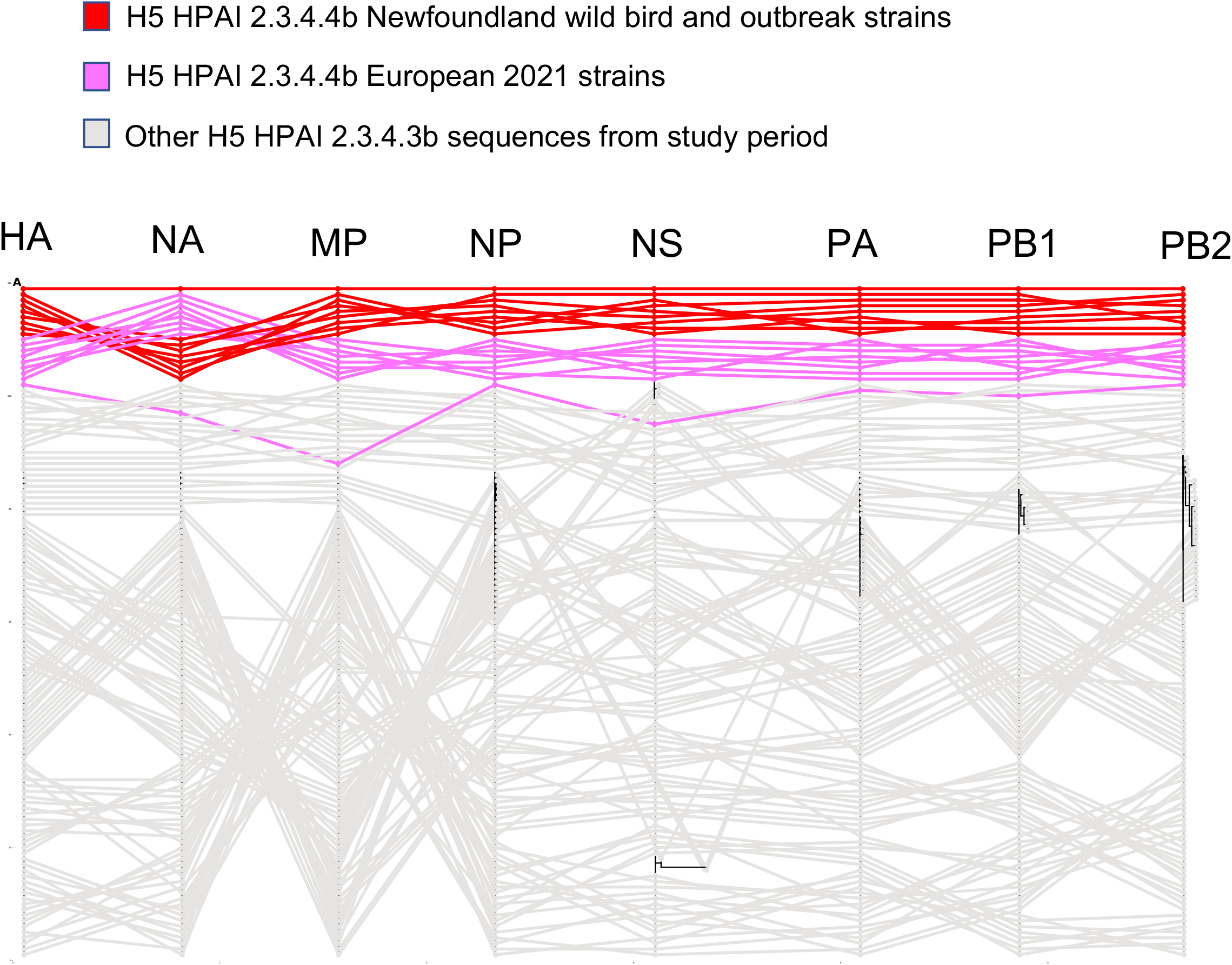
(because of its large size supplementary figure 2 is submitted as related document) Phylogenetic incongruence analyses. Maximum likelihood trees for the H and N gene segments and internal gene segments from equivalent strains were connected across the trees. Tips and connecting lines are coloured according to the legend.

**Table S1.** Common and Latin species names of the birds mentioned in the text.

| Common name | Scientific name |
| --- | --- |
| Atlantic puffin | *Fratercula arctica* |
| Black-legged kittiwake | *Rissa tridactyla* |
| Barnacle goose | *Branta leucopsis* |
| Black-headed gull | *Chroicocephalus ridibundus* |
| Brunnich's guillemot | *Uria lomvia* |
| Common eider | *Somateria mollissima* |
| Common starling | *Sturnus vulgaris* |
| Eurasian teal | *Anas crecca* |
| Eurasian wigeon | *Mareca penelope* |
| Feral pigeon | *Columba livia domestica* |
| Great black-backed gull | *Larus marinus* |
| Great skua | *Stercorarius skua* |
| Greater white-fronted goose | *Anser albifrons* |
| Greylag goose | *Anser anser* |
| Lesser black-backed gull | *Larus fuscus* |
| Light-bellied brent goose | *Branta bernicla* |
| Mallard | *Anas platyrhynchos* |
| Northern fulmar | *Fulmarus glacialis* |
| Northern pintail | *Anas acuta* |
| Pink-footed goose | *Anser brachyrhynchus* |
| Purple sandpiper | *Calidris maritima* |
| Red knot | *Calidris canutus* |
| Snow goose | *Anser caerulescens* |
| Snowy egret | *Egretta thula* |
| Tufted duck | *Aythya fuligula* |
| Whooper swan | *Cygnus cygnus* |

**Table S2.** (because of its large size supplementary table 2 is submitted as related document) **Ring-recoveries in Newfoundland of species most frequently encountered.** Data on transatlantic migration of wild bird species, including wintering, staging and breeding grounds, population sizes, numbers of birds encountered on Newfoundland and their breeding origin, number of ring recoveries for various transects of bird migration routes and timing of spring and autumn migration. Names are marked in bold for species that were found to be HPAI-H5-positive between October 2020 and April 2021. For the numbers of birds observed in Newfoundland, first the annual average between 2011 - 2021 is given followed by the number of birds in the autumn of 2021, unless otherwise stated. When exact numbers are unknown, estimated number of recoveries is marked as + (1-10), ++ (10-100) or +++ (100-1000).

**Table S3.** (because of its large size supplementary table 3 is submitted as related document) HPAI-H5-positive wild bird species by country and month, from October 2020 to December 2021 (based on OIE, IZSVENEZIE, and APHA online reports).

**Table S4.** List of samples for virological analysis from different farm animals.

| Bird type | Number on site | Number of oropharyngeal swabs | Number of cloacal swabs | Number of sera |
| --- | --- | --- | --- | --- |
| Turkey<br>(*Meleagris gallopavo*) | 4 | 4 | 4 | 4 |
| Silkie chicken<br>(*Gallus gallus*) | 8 | 8 | 8 | 0 |
| Emus<br>(*Dromaius novaehollandiae*) | 2 | 2 | 0 | 0 |
| Peafowl<br>(*Pavo cristatus*) | 5 | 5 | 5 | 0 |
| Domestic goose<br>(*Anser anser*) | 14 | 0 | 14 | 5 |
| Domestic duck<br>(*Anas platyrhynchos domesticus*) | 20 | 0 | 20 | 7 |
| Guineafowl<br>(*Numida meleagris*) | 0 | 0 | 0 | 0 |
| Chicken-other<br>(*Gallus gallus*) | 4 | 4 | 4 | 1 |
| Total | 57 | 23 | 55 | 17 |

## References

1. L. Duan, J. Bahl, G. J. Smith, J. Wang, D. Vijaykrishna, L. I. Zhang, J. X. Zhang, K. S. Li, X. H. Fan, C. L. Cheung, K. Huang, L. L. Poon, K. F. Shortridge, R. G. Webster, J. S. Peiris, H. Chen, Y. Guan, The development and genetic diversity of H5N1 influenza virus in China, 1996-2006. Virology. 380, 243–54 (2008). doi: 10.1016/j.virol.2008.07.038.

2. Global Consortium for H5N8 and Related Influenza Viruses. Role for migratory wild birds in the global spread of avian influenza H5N8. Science. 354, 213–217 (2016). doi: 10.1126/science.aaf8852.

3. S. J. Lycett, A. Pohlmann, C. Staubach, V. Caliendo, M. Woolhouse, M. Beer, T. Kuiken; Global Consortium for H5N8 and Related Influenza Viruses. Genesis and spread of multiple reassortants during the 2016/2017 H5 avian influenza epidemic in Eurasia. Proc Natl Acad Sci U S A. 117, 20814–20825 (2020). doi: 10.1073/pnas.2001813117.

4. EFSA (European Food Safety Authority), ECDC (European Centre for Disease Prevention and Control), EURL (European Reference Laboratory for Avian Influenza), C. Adlhoch, A. Fusaro, J. L. Gonzales, T. Kuiken, S. Marangon, E. Niqueux, C. Staubach, C. Terregino, I. Aznar, I. Muñoz Guajardo, F. Baldinelli, EFSA Scientific report: Avian influenza overview September-_December 2021. 94 pp. (2021). https://doi.org/10.2903/j.efsa.2021.7108.

5. B. Olsen, V.J. Munster, A. Wallensten, J. Waldenström, A. D. Osterhaus, R. A. Fouchier, Global patterns of influenza a virus in wild birds. Science. 312, 384–8 (2006). doi: 10.1126/science.1122438.

6. C. Adlhoch, F. Baldinelli, A. Fusaro, C. Terregino, Avian influenza, a new threat to public health in Europe? Clin Microbiol Infect. 2021 Nov 9:S1198-743X(21)00632-7 (2021). doi: 10.1016/j.cmi.2021.11.005.

7. W. Xu, Y. Berhane, C. Dubé, B. Liang, J. Pasick, G. VanDomselaar, S. Alexandersen, Epidemiological and Evolutionary Inference of the Transmission Network of the 2014 Highly Pathogenic Avian Influenza H5N2 Outbreak in British Columbia, Canada. Sci Rep. 6:30858 (2016). doi: 10.1038/srep30858.

8. OIE 2021, report id. CAN-2021-HPAI-001, https://wahis.oie.int/#/report-info?reportId=45054 (Accessed 08/01/2022).

9. F. Di Giallonardo, J.L. Geoghegan, D.E. Docherty, R.G. McLean, M.C. Zody, J. Qu, X. Yang, B.W. Birren, C.M. Malboeuf, R.M. Newman, H.S. Ip, E.C. Holmes, Fluid Spatial Dynamics of West Nile Virus in the United States: Rapid Spread in a Permissive Host Environment. Journal of virology, 90, 862–872 (2015). https://doi.org/10.1128/JVI.02305-15.

10. R. J. Dusek, G. T. Hallgrimsson, H. S. Ip, H, J. E. Jónsson, S. Sreevatsan, S. W. Nashold, J. L. TeSlaa, S. Enomoto, R. A. Halpin, X. Lin, N. Fedorova, T. B. Stockwell, V. G. Dugan, D. E. Wentworth, J. S. Hall, North Atlantic migratory bird flyways provide routes for intercontinental movement of avian influenza viruses. PloS one, 9, e92075 (2014). https://doi.org/10.1371/journal.pone.0092075.

11. J. S. Hall, G. T. Hallgrimsson, K. Suwannanarn, S. Sreevatsen, H. S. Ip, E. Magnusdottir, J. L. TeSlaa, S. W. Nashold, R. J. Dusek, Avian influenza virus ecology in Iceland shorebirds: Intercontinental reassortment and movement. Infection, Genetics and Evolution, 28, 130–136 (2014). https://doi.org/10.1016/j.meegid.2014.09.013.

12. A. D. Fox, J. O. Leafloor, (eds.), A Global Audit of the Status and Trends of Arctic and Northern Hemisphere Goose Populations (Component 2: Population accounts). Conservation of Arctic Flora and Fauna International Secretariat: Akureyri, Iceland. ISBN 978-9935-431-74-5 (2018).

13. eBird. 2022. eBird: An online database of bird distribution and abundance [web application]. eBird, Cornell Lab of Ornithology, Ithaca, New York. Available: http://www.ebird.org. (Accessed: Date January 4, 2022)

14. S. N. G. Howell, I. Lewington, W. Russell, (eds.), Rare birds of North America. Princeton University Press, USA (2014).

15. M. C. Edgell, Trans-hemispheric movements of Holarctic Anatidae: the Eurasian wigeon (Anas penelope L.) in North America. Journal of biogeography, 11, 27–39 (1984).

16. M. L. Mallory, A. J. Fontaine, (eds.) Key marine habitat sites for migratory birds in Nunavut and the Northwest Territories. Canadian Wildlife Series Occasional Paper Number 109. Environment Canada. Ottawa, Canada (2004).

17. A. J. Gaston, M. L. Mallory, H. G. Gilchrist, Population and trends of Canadian Arctic seabirds. Polar Research, 35, 1221–1232 (2012).

18. D. A. Fifield, A. Hedd, S. Avery-Gomm, G. J. Robertson, C. Gjerdrum, L. McFarlane Tranquilla, Employing predictive spatial models to inform conservation planning for seabirds in the Labrador Sea. Frontiers in Marine Science 4:149 (2017). doi: 10.3389/fmars.2017.00149.

19. S. N. P. Wong, C. Gjerdrum, K. H. Morgan, and M. L. Mallory, Hotspots in cold seas: The composition, distribution, and abundance of marine birds in the North American Arctic, J. Geophys. Res. Oceans, 119, 1691–1705 (2014).

20. K. Kuletz, M. Mallory, G. Gilchrist, G. Robertson, F. Merkel, B. Olsen, E. Hansen, M. Rönkä, T. Anker-Nilssen, H. Strøm, S. Déscamps, M. Gavrilo, R. Kaler, D. Irons, A. Below. Seabirds. Pages 139-165 in State of the arctic marine biodiversity report. Conservation of Arctic Flora and Fauna, Circumpolar Biodiversity Monitoring Plan, Arctic Council. Reykjavik, Iceland (2017).

21. A. D. Fox, C. Glahder, C. R. Mitchell, D. A. Stroud, H. Boyd, J. Frikke, North American Canada Geese (Branta canadensis) in West Greenland. The Auk, 131, 231–233 (1996).

22. D.F. Sherony, Greenland geese in North America. Birding, 40, 46–56 (2008).

23. J. van den Brand, J.H. Verhagen, E. Veldhuis Kroeze, M. van de Bildt, R. Bodewes, S. Herfst, M. Richard, P. Lexmond, T.M. Bestebroer, R. Fouchier, T. Kuiken, (2018). Wild ducks excrete highly pathogenic avian influenza virus H5N8 (2014-2015) without clinical or pathological evidence of disease. Emerging microbes & infections, 7, 67 (2018). https://doi.org/10.1038/s41426-018-0070-9.

24. L. M. Tuck, The occurrence of Greenland and European birds in Newfoundland. Bird-Banding. 42, 84–209 (1971).

25. M. Frederiksen, B. Moe, F. Daunt, R. A. Phillips, R. T. Barrett, M. I. Bogdanova, T. Boulinier, J. W. Chardine, O. Chastel, L. S. Chivers, S. Christensen-Dalsgaard, C. Clément-Chastel, K. Colhoun, R. Freeman, A. J. Gaston, J. González-Solís, A. Goutte, D. Grémillet, T. Guilford, G. H. Jensen, Y. Krasnov, S. H. Lorentsen, M. Mallory, M. Newell, B. Olsen, D. Shaw, H. Steen, H. Strøm, G. H. Systad, T. L. Thórarinsson, T. Anker-Nilssen, Multicolony tracking reveals the winter distribution of a pelagic seabird on an ocean basin scale. Diversity and Distributions, 18, 530–542 (2012).

26. T. E. Davies, A. P. B. Carneiro, M. Tarzia, E. Wakefield, J. C. Hennicke, M. Frederiksen, E. S. Hansen, B. Campos, C. Hazin, B. Lascelles, B, Multispecies tracking reveals a major seabird hotspot in the North Atlantic. Conservation Letters, 14, e12824 (2021). https://doi.org/10.1111/conl.12824.

27. M. Wille, G. J. Robertson, H. Whitney, D. Ojkic, A. S. Lang, Reassortment of American and Eurasian genes in an influenza A virus isolated from a great black-backed gull (Larus marinus), a species demonstrated to move between these regions. Arch. Virol. 156, 107–115 (2011). https://doi.org/10.1007/s00705-010-0839-1.

28. A. S. Lang, C. Lebarbenchon, A. M. Ramey, G. J. Robertson, J. Waldenström, M. Wille, Assessing the Role of Seabirds in the Ecology of Influenza A Viruses. Avian Dis. 60, 378–86 (2016). doi: 10.1637/11135-050815-RegR.

29. P. Sagulenko, V. Puller, R. A. Neher, TreeTime: Maximum-likelihood phylodynamic analysis. Virus Evol. 4, vex042 (2018). doi:10.1093/ve/vex042

30. M. J. Poen, D. Venkatesh, T. M. Bestebroer, O. Vuong, R. D. Scheuer, B. B. Oude Munnink, D. de Meulder, M. Richard, T. Kuiken, M. P. G. Koopmans, L. Kelder, Y. J. Kim, Y. J. Lee, M. Steensels, B. Lambrecht, A. Dan, A. Pohlmann, M. Beer, V. Savic, I. H. Brown, R. A. M. Fouchier, N. S. Lewis, Co-circulation of genetically distinct highly pathogenic avian influenza A clade 2.3.4.4 (H5N6) viruses in wild waterfowl and poultry in Europe and East Asia, 2017-18. Virus Evol. 5, vez004 (2019). doi: 10.1093/ve/vez004.

31. S. M. Billerman, B. K. Keeney, P. G. Rodewald, and T. S. Schulenberg (eds). Birds of the World. Cornell Laboratory of Ornithology, Ithaca, NY, USA. (2020) https://birdsoftheworld.org/bow/home.

32. A.D. Fox, J.O. Leafloor, (eds.). A Global Audit of the Status and Trends of Arctic and Northern Hemisphere Goose Populations (Component 2: Population accounts). Conservation of Arctic Flora and Fauna International Secretariat: Akureyri, Iceland. ISBN 978-9935-431-74-5 (2018).

33. Icelandic Institute of Natural History, 2021, https://en.ni.is/biota/animalia/chordata/aves, accessed 30/12/2021.

34. M. van Roomen, C. van Turnhout, J. Blew, K. Koffijberg, S. Nagy, G. Citegetse, R. Foppen, East Atlantic Flyway. In: Wadden Sea Quality Status Report 2017. Eds.: Kloepper S. et al., Common Wadden Sea Secretariat, Wilhelmshaven, Germany (2018).

35. K. Brides, K.A. Wood, C. Hall, B. Burke, G. McElwaine, O. Einarsson, E.C. Rees, The Icelandic Whooper Swan Cygnus cygnus population: current status and long-term (1986-2020) trends in its numbers and distribution. Wildfowl, 71, 29–57 (2021).

36. A.D. Fox, D. Sinnett, J.A. Baroch, D.A. Stroud, K. Kampp, C. Egevang, D. Boertmann, (eds.). The status of Canada Goose Branta canadensis subspecies in Greenland (2012).

37. A. Mosbech, G. Gilchrist, F. Merkel, C. Sonne, A. Flagstad, H. Nyegaard, Year-round movements of Northern Common Eiders Somateria mollissima borealis breeding in Arctic Canada and West Greenland followed by satellite telemetry. ARDEA-WAGENINGEN-, 94, 651 (2006).

38. E. Magnusdottir, E.H. Leat, S. Bourgeon, H. Strøm, A. Petersen, R.A. Phillips, S.A. Hanssen, J.O. Bustnes, P. Hersteinsson, R.W. Furness, Wintering areas of great skuas Stercorarius skua breeding in Scotland, Iceland and Norway. Bird Study, 59, pp.1–9 (2012).

39. D. Boertmann, The lesser black-backed gull, Larus fuscus, in Greenland. Arctic, pp.129–133 (2008).

40. G.J. Robertson, Current status of the Manx Shearwater (Puffinus puffinus) colony on Middle Lawn Island, Newfoundland. Northeastern Naturalist, 9, pp.317–324 (2002).

41. P. Fauchald,, A. Tarroux, V.S. Bråthen, S. Descamps, M. Ekker, H.H. Helgason, B. Merkel, B. Moe, J. Åström, H. Strøm, Arctic-breeding seabirds’ hotspots in space and time a methodological framework for year-round modelling of abundance and environmen-tal niche using light-logger data. NINA Report 1657. Norwegian Institute for Nature Re-search (2019).

42. R.L. Stirnemann, J.O.H.N. O’Halloran, M. Ridgway, A. Donnelly, Temperature-related increases in grass growth and greater competition for food drive earlier migrational departure of wintering Whooper Swans. Ibis, 154, 542–553 (2012).

43. G.T. Hallgrimsson, H.V. Gunnarsson, O. Torfason, R.J. Buijs, K.C. Camphuysen, Migration pattern of Icelandic Lesser Black-backed Gulls Larus fuscus graellsii: indications of a leap-frog system. Journal of Ornithology, 153, 603–609 (2012).

44. B. Merkel, S. Descamps, N.G. Yoccoz, J. Danielsen, F. Daunt, K.E. Erikstad, H. Strøm (2019). Earlier colony arrival but no trend in hatching timing in two congeneric seabirds (Uria spp.) across the North Atlantic. Biology letters, 15, 20190634 (2019).

45. K. Brides, K.A. Wood, C. Hall, B. Burke, G. McElwaine, O. Einarsson, E.C. Rees, The Icelandic Whooper Swan Cygnus cygnus population: current status and long-term (1986–2020) trends in its numbers and distribution. Wildfowl, 71, 29–57 (2021).

46. B. Swann, I.K. Brockway, M. Frederiksen, R.D. Hearn, C. Mitchell, A/ Sigfússon, (2005) Within-winter movements and site fidelity of Icelandic Greylag Geese Anser anser. Bird Study, 52, 25–36 (2005).

47. G.T. Hallgrimsson, H.V. Gunnarsson, O. Torfason, R.J. Buijs, K.C. Camphuysen, Migration pattern of Icelandic Lesser Black-backed Gulls Larus fuscus graellsii: indications of a leap-frog system. Journal of Ornithology, 153, 603–609 (2012).

48. Canadian Wildlife Service Waterfowl Committee. Population Status of Migratory Game Birds in Canada. November 2019. CWS Migratory Birds Regulatory Report Number 52 (2020).

49. D. A. Fifield, K. P. Lewis, C. Gjerdrum, G. J. Robertson, R. Wells. Offshore Seabird Monitoring Program. Environment Studies Research Funds Report No. 183. St. John’s. 68 p. (2009).

50. P. Lyngs, Migration and winter ranges of birds in Greenland. Dansk Ornitologisk Forenings Tidsskrift,97, 1–167 (2003).

